# Erbin interacts with NHERF1 and Ezrin to stabilize a membrane ErbB2 signaling complex in HER2-positive breast cancer

**DOI:** 10.1101/2024.10.01.616146

**Authors:** Jaekwang Jeong, Kwangmin Yoo, Jongwon Lee, Jae Hun Shin, Jungmin Choi, John Wysolmerski

## Abstract

Approximately 20% of breast cancers overexpress ErbB2/HER2/Neu, a receptor tyrosine kinase. Our previous studies demonstrated that HER2 interacts with the calcium pump, PMCA2, and the scaffolding molecules, NHERF1 and Ezrin to stabilize HER2/HSP90 interactions and contribute to the retention of active HER2 at the plasma membrane. In the normal mammary epithelium where apical/basal polarity is tightly regulated by junctional proteins, HER2 is expressed at the basolateral membrane and interacts with the LAP family member, Erbin, whereas PMCA2, NHERF1, and Ezrin localize to the apical membrane. Here, we show that loss of apical membrane polarity in hyperplastic lesions of MMTV-Neu mammary glands or in human DCIS leads to intermixing of these molecules and allows Erbin to interact with NHERF1, Ezrin and HER2 initially within the basolateral membrane and then more diffusely within the plasma membrane. In SKBR3 cells, Erbin physically interacts with NHERF1, Ezrin and HER2 in actin-rich membrane protrusions that we have previously described to be sites of active HER2 signaling. Knockdown of Erbin in these cells reduced HER2 signaling by disrupting the formation of a HER2/NHERF1/Ezrin/HSP90 protein complex in the membrane protrusions. Furthermore, inhibition of Ezrin or knock-down of NHERF1 expression disrupted the ability of Erbin to interact with HER2. Taken together, our data suggest that Erbin supports HER2 stability, HER2 membrane retention and HER2 transforming ability by interacting with Ezrin and NHERF1 to maintain a multi-protein signaling complex necessary for HER2-mediated transformation.

## Introduction

Approximately 20% of breast cancers overexpress the receptor tyrosine kinase, ERBB2/HER2/Neu (HER2), usually through amplification of the *ERBB2* gene. MMTV-Neu transgenic mice overexpress HER2 in mammary epithelial cells which results in the development of mammary tumors, demonstrating that HER2 overexpression is directly oncogenic in the breast (1, 2). Overexpression of HER2 leads to more aggressive tumor behavior and increased mortality (3–5), although this has been partially mitigated by the development of targeted therapies directed at HER2 protein. However, as with other targeted therapies, many patients develop resistance and stop responding to current anti-HER2 therapy, underscoring the ongoing need for more and better therapies for HER2-positive cancers. Unlike other ErbB family members, activated HER2 tends to remain on the cell surface and signal for prolonged periods, a characteristic that contributes to its transforming ability. Previous studies from our lab and others have demonstrated that interactions between HER2 and the scaffolding molecules, NHERF1 and Ezrin, stabilize interactions between HER2 and HSP90, MAL2 and the calcium pump, PMCA2, all of which contribute to the plasma membrane retention of HER2 within lipid raft- and actin-rich membrane domains that protrude from the surface of cancer cells. These proteins interact via PDZ domains and the formation of this multiprotein signaling complex is required for prolonged downstream biochemical signaling that drives cell transformation (6–8).

Interestingly, a higher percentage (40% instead of 20%) of ductal carcinoma in situ (DCIS) lesions than invasive carcinomas demonstrate HER2 overexpression (9). The underlying mechanism for this higher proportion of HER2-positive DCIS and its significance remain unclear but this observation suggests that upregulation of HER2 may play a significant role during the early transformation of a significant proportion of breast cancers. In cell lines, it has been shown that increased expression of HER2 disrupts cell polarity and adhesion (10), important characteristics of cancer cells (11, 12). Disruption of apical-basal polarity is mediated by interactions of HER2 with PAR6 and aPKC, which, in turn, disrupt their interactions with Par3 (12). Therefore, HER2 overexpression/signaling contributes to transformation of breast cells, in part, by disrupting the Par complex that maintains apical-basolateral compartmentalization within normal mammary epithelial cells.

Erbin is a cytoplasmic scaffolding protein that contains both a leucine-rich repeat (LRR) and a PSD95/Dlg1/zo1 (PDZ) domain (13, 14). It was initially described due to its ability to specifically bind ErbB2, but it does not bind to other ErbB proteins (15, 16). Erbin interacts with HER2 at the basolateral membrane of normal epithelial cells via its PDZ domain (16). These interactions have been shown to be necessary for HER2 to bind HSP90 and contribute to the ability of HER2 to transform mammary epithelial cells in vitro as well as in MMTV-Neu mice *in vivo*. Our previous studies identified very similar influences of PMCA2, NHERF1 and Ezrin on HER2 signaling and/or its transforming ability (6–8) (15). Therefore, in this study we examined whether Erbin interacts with the previously defined HER2/NHERF1/Ezrin protein complex. Our findings suggest that Erbin is required for interactions between HER2, NHERF1, Ezrin and HSP90 that stabilize active HER2 at the plasma membrane after disruption of apical polarity allows intermixing of apical and basolateral compartments.

## Results

### Colocalization of HER2 with NHERF1 and Ezrin during the onset of HER2-driven tumorigenesis

In nulliparous and lactating female mice, luminal mammary epithelial cells express Ezrin and NHERF1 at the apical plasma membrane (Fig. 1A) (8, 17). We have not been able to detect ErbB2/HER2 expression in normal mammary epithelial cells by immunofluorescence, but when we examined HER2-overexpressing, MMTV-neu mice prior to tumor formation, we detect HER2 expression within the basolateral membrane (Fig. 1B). However, in lactating MMTV-neu mice, Ezrin and NHERF1 remain localized to the apical membrane (Fig. 1B). In contrast, in areas of alveolar hyperplasia in MMTV-Neu mice, HER2 expression begins to co-localize with ezrin and NHERF1 more generally around the plasma membrane of the cells, representing a breakdown of their normally polarized patterns (Fig. 1C). These observations prompted us to examine the expression of these three molecules in early human breast cancer samples. We analyzed the localization of HER2, NHERF1, and Ezrin in HER2-positive human ductal carcinoma in situ (DCIS). As with hyperplastic lesions in mice, lumen filling DCIS samples had disrupted apical-basal polarity as evidenced by the generalized expression of the sodium-potassium ATPase throughout the plasma membrane (Fig.1D). In this context, HER2 co-localized with NHERF1 and Ezrin within the entire plasma membrane (Fig. 1D). Interestingly, in some sections we noted increased HER2 expression within what appeared, histologically, to be normal luminal epithelial cells at the periphery of DCIS lesions (Fig. 1E). In these areas, Na/ATPase expression remained restricted to the basolateral membrane and colocalized with HER2 expression (Fig. 1E). In contrast, NHERF1 and Ezrin expression could be detected in both apical and basolateral membranes, where they co-localized with HER2 expression (Fig. 1E), consistent with prior reports that loss of apical polarity is an early event in HER2-mediated transformation of human mammary epithelial cells *in vitro* (12, 18). We next utilized published RNAseq data to examine levels of gene expression of ErbB2 (HER2), NHERF1 and Ezrin in HER2-positive human DCIS as compared to levels of expression of the same mRNAs in normal breast tissue. Gene set enrichment analysis (GSEA) showed the deactivation of “hallmark apical junction” pathway in DCIS consistent with a loss of apical polarity (Fig. 1F&G). In normal tissue, there was no significant correlation between ErbB2 and NHERF1 expression or between ErbB2 and Ezrin expression (Fig. 1H). In contrast, we found a positive correlation between Ezrin and ErbB2 expression in DCIS, although there was no correlation between NHERF1 and ErbB2 expression (Fig. 1H). We also examined the spatial correlation between the expression of ErbB2 (HER2), SLC9A3R1 (NHERF1), and EZR (Ezrin) in breast cancer cells, using a spatial transcriptomics (Visium) database of human breast cancer (19). Figure 2A-C shows a spatial profile of relative gene expression of ErbB2, SLC9A3R1, and EZR. Non-malignant cells, including adipocytes, immune and stromal cells, and myoepithelial cells were chosen as a negative control since normal luminal epithelial cells were not present in this tissue section (20). ErbB2 and NHERF1 transcript levels were higher in DCIS and invasive tumor cells than in non-malignant and myoepithelial cells (Fig. 2A&C). Interestingly, Ezrin expression was significantly higher only in DCIS cells but not in the invasive tumor cells as compared to non-malignant and myoepithelial cells (Fig. 2A&C). Then, DCIS spots were divided by SLC9A3R1/EZR high and low according to median (Fig. 2D). Gene set enrichment analysis (GSEA) showed the deactivation of “hallmark apical junction” pathway in SLC9A3R1/EZR^high^ DCIS compared to SLC9A3R1/EZR^low^ DCIS (Fig. 2E&F).

**Figure 1.**
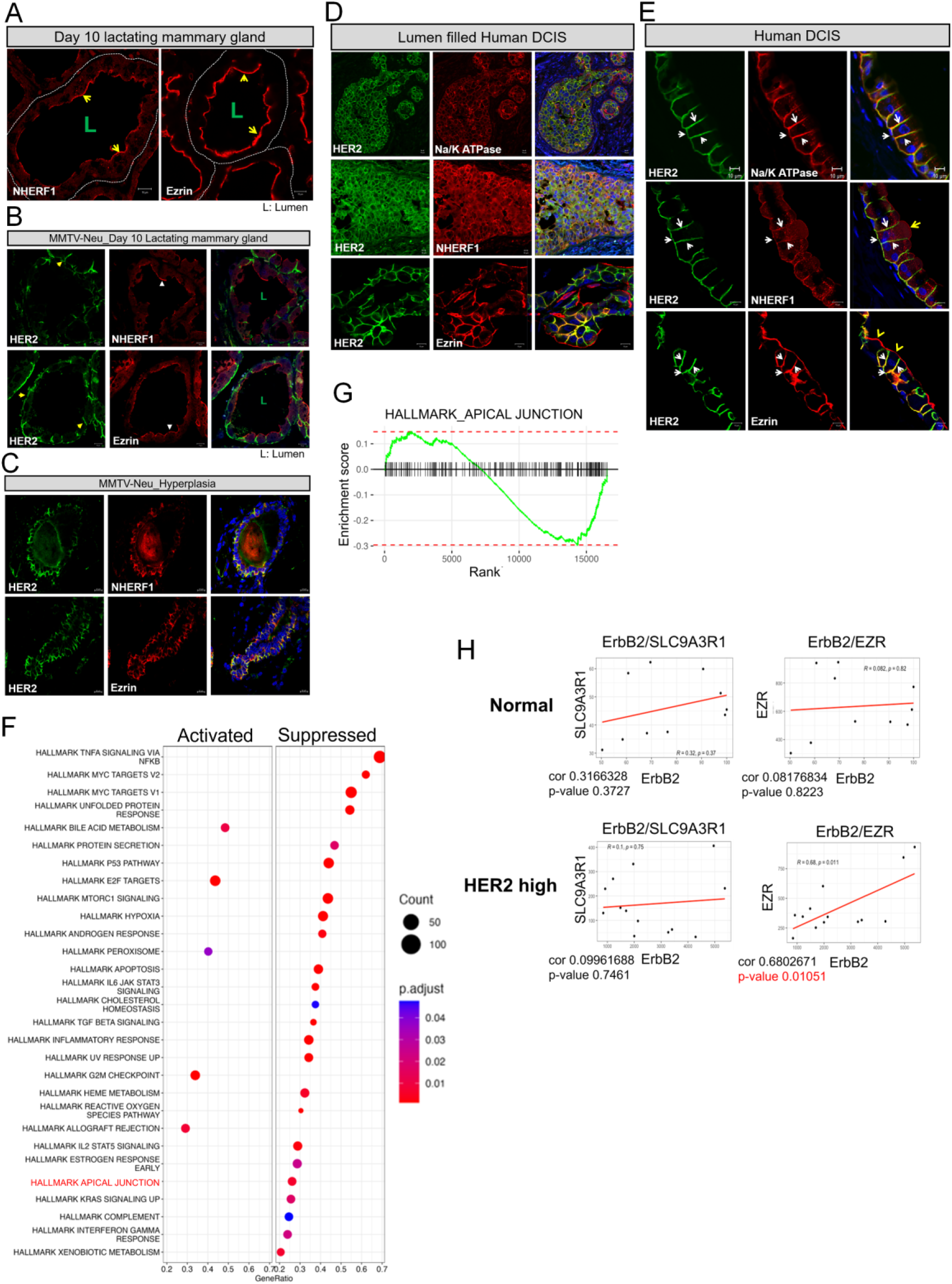
Association of HER2, NHERF1, and Ezrin in hyperplasia and DCIS. A) Immunofluorescence for NHERF1 and Ezrin in day 10 lactating WT mouse mammary glands. All scale bars represent 10μM. B) Immunofluorescence for HER2 with NHERF1 and Ezrin in day 10 lactating MMTV-Neu mouse mammary glands. All scale bars represent 10μM. C) Immunofluorescence for HER2 with NHERF1 (top row), and Ezrin (bottom row) in hyperplastic lesions from MMTV-Neu mammary glands. All scale bars represent 10μM. D) Immunofluorescence for HER2 with Na/K ATPase (top row), NHERF1 (middle row), and Ezrin (bottom row) in human lumen filled DCIS sections. All scale bars represent 10μM. E) Immunofluorescence for HER2 with Na/K ATPase (top row), NHERF1 (middle row), and Ezrin (bottom row) in human sections adjacent to DCIS lesions. All scale bars represent 10μM. F) Gene Set Enrichment Analysis (GSEA) from RNAseq comparing tumor free human breast tissue (normal) vs human HER2 positive DCIS. G) Enrichment score of apical_junction pathway for HER2 positive DCIS from Hallmark GSEA analysis. H) Correlation of HER2 gene expression with SLC9A3R1 (NHERF1) and EZR (Ezrin) from RNAseq comparing normal breast vs human HER2 positive DCIS.

**Figure. 2.**
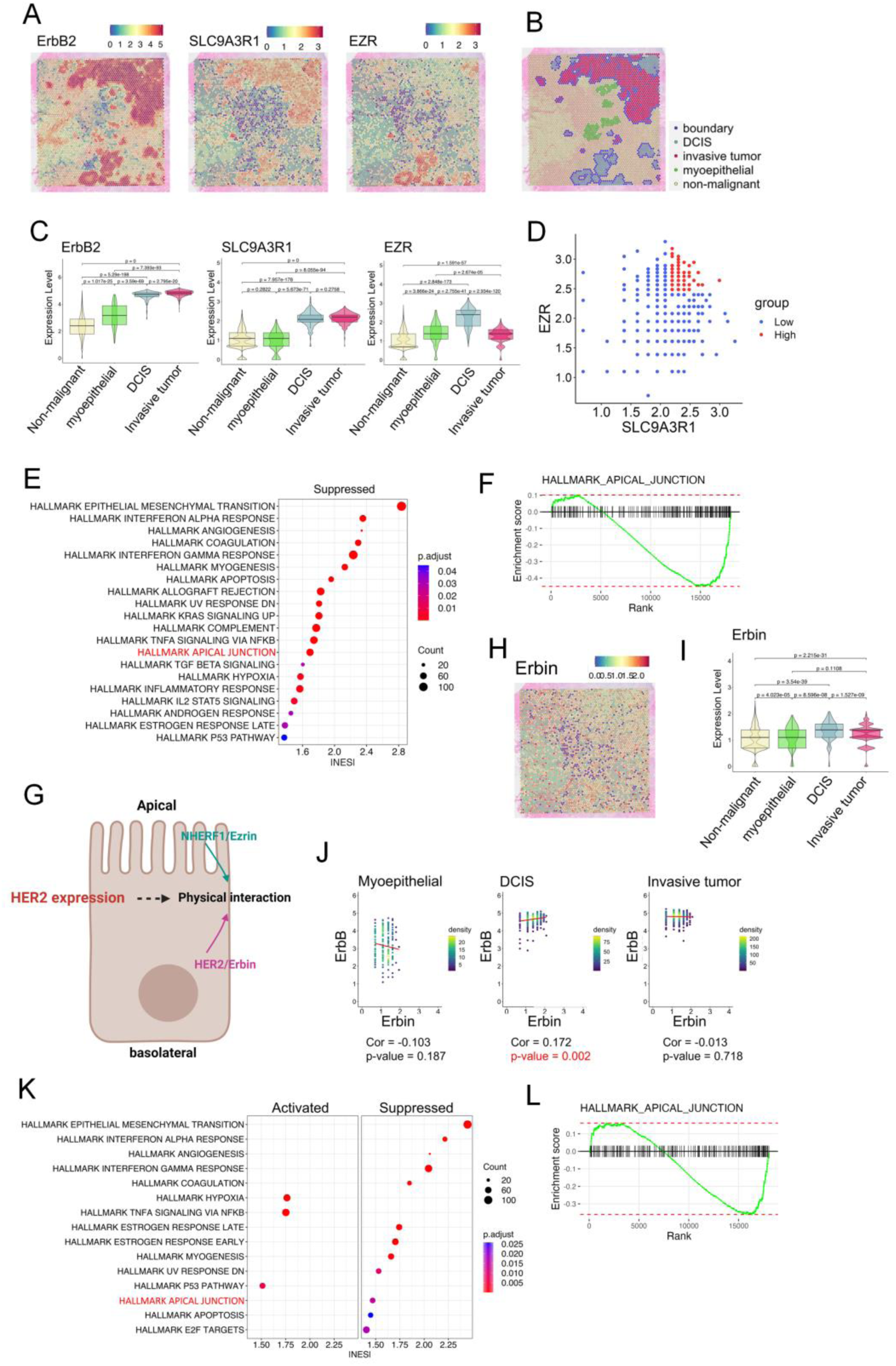
Co-expression of Erbin with HER2, SLC9A3R1 (NHERF1), and EZR (Ezrin). A) Cell type specific marker genes [ERBB2 (HER2), SLC9A3R1 (NHERF1), and EZR (Ezrin)] are expressed as log_2_ (normalized UMI counts). B) Spatial distribution of cells identified by unsupervised clustering based on differential gene expression analysis. C) Violin plot illustrating the selected marker genes expression of each cell type [ERBB2 (HER2), SLC9A3R1 (NHERF1), and EZR (Ezrin)]. The center line in each box depict the medians, and the top and bottom edges depict the first/third quartiles. D) Diagram show separated spots of SLC9A3R1/EZR^high^ (red) and SLC9A3R1/EZR^LOW^ (blue) in DCIS. E) Gene Set Enrichment Analysis (GSEA) from Visium comparing SLC9A3R1/EZR^high^ and SLC9A3R1/EZR^LOW^ in DCIS. F) GSEA enrichment score of apical_junction pathway from SLC9A3R1/EZR^high^ vs. SLC9A3R1/EZR^LOW^ in DCIS. G) Diagram of proposed pathway by which HER2 expression could allow interaction between HER2/Erbin and NHERF1/Ezrin complexes. Created with BioRender.com. H) Erbin expression as log_2_ (normalized UMI counts). I) Violin plot illustrating Erbin expression in each cell type. J) Pearson correlation coefficient between Erbin and ERBB2 in each cell type. K) Gene Set Enrichment Analysis (GSEA) from Visium comparing Erbin^high^ and Erbin^LOW^ in DCIS. F) GSEA enrichment score of hallmark apical_junction pathway from the comparison between Erbin^high^ and Erbin^LOW^ DCIS.

Erbin is a scaffolding molecule previously shown to a interact with HER2 and to influence cellular proliferation and tumorigenesis in the MMTV-neu transgenic mouse model of HER2 overexpressing breast cancer (Fig. 2G) (16, 21). We previously demonstrated physical interactions between NHERF1, Ezrin and HER2 that require the PDZ domains of NHERF1 and that facilitate HER2 driven tumorigenesis (7, 8). Like NHERF1, Erbin also contains a PDZ domain implying the possibility of physical interactions between Erbin and the NHERF1/Ezrin/HER2 complex (Fig. 2G). We first examined Erbin gene expression using the aforementioned spatial transcriptomics (Visium) data derived from human breast cancer (19). We found significant increase in Erbin expression in DCIS and invasive tumor areas as compared to non-malignant and myoepithelial cells, although the absolute increase appeared greater for DCIS than for invasive tumor (Fig. 2H-I). Furthermore, when we examined correlations between the expression of Erbin vs. these other genes, we found a statistically significant positive correlation between Erbin expression and ErbB2 expression in DCIS cells but not in invasive tumor cells (Fig. 2J). Interestingly, GSEA showed the deactivation of “hallmark apical junction” pathway in Erbin^high^ DCIS when compared to Erbin^low^ DCIS (Fig. 2K-L). Together, these data demonstrate correlations between ErbB2, NHERF1 and Ezrin gene expression as well as the colocalization of ErbB2, NHERF1, and Ezrin proteins early during the development of HER2-positive breast cancer.

### Erbin interacts with the HER2/NHERF1/Ezrin protein complex

Next, we tested whether there were protein-protein interactions between the same molecules during HER2-driven tumorigenesis. In order to examine potential colocalization of Erbin with HER2 in non-transformed mammary epithelial cells, we examined WT mice and MMTV-Neu mice prior to tumor development. As shown in Fig. 3A, in lactating WT mice, we saw low level cytoplasmic immunofluorescence staining and some apparent nuclear staining. However, in lactating MMTV-Neu mice, Erbin immunofluorescence was more evident and appeared both in the cytoplasm but also became more restricted to the basolateral membrane, where it co-localized with ErbB2 (Fig. 3A). In contrast to normal mammary epithelial cells, in hyperplastic lesions from MMTV-Neu mice, Erbin colocalized with HER2, NHERF1, and Ezrin in the plasma membrane either more uniformly around both the apical and basolateral membranes or more concentrated within the basolateral memebrane (Fig. 3B). Likewise, in HER2 positive human samples Erbin also co-localized with HER2, NHERF1 and Ezrin generally throughout the plasma membrane (Fig. 3C). This was also true in SKBR3 cells, a HER2-positive breast cancer cell line. As shown in Fig. 3D, Erbin co-localized with HER2, NHERF1, and Ezrin at the plasma membrane and especially within membrane protrusions, which we have previously documented to be an important location of active HER2 signaling in breast cancer cells. Interestingly, erbin did not co-localize with NHERF1 or ezrin in MCF10A cells, a non-transformed mammary epithelial cell line that does not express HER2 (Fig. 3D-E) (6–8, 22, 23). Consistent with the immunofluorescence data, ezrin and erbin were pulled down when HER2 was immunoprecipitated from SKBR3 cells (Fig. 3F). We have not been able to IP NHERF1 with available antibodies, but using an anti-HA antibody, we were able to co-IP Erbin with NHERF1 in SKBR3 cells transiently expressing an HA-tagged NHERF1, demonstrating physical interactions between Erbin and NHERF1 as well, but not in MCF10A cells (Fig. 3G). Finally, we used proximity ligation assays (PLA) to demonstrate that Erbin is in close physical proximity to HER2, Ezrin and NHERF1 in SKBR3 cells (Fig. 3H). However, again, interactions between HER2, NHERF1, Erzin and Erbin did not occur in MCF10A cells that don’t express HER2. Together, these findings suggest that expression of HER2 and loss of cell polarity allows physical interactions between Erbin, NHERF1, Ezrin and HER2 in breast cancer cells that do not normally occur in non-malignant mammary epithelial cells.

**Figure 3.**
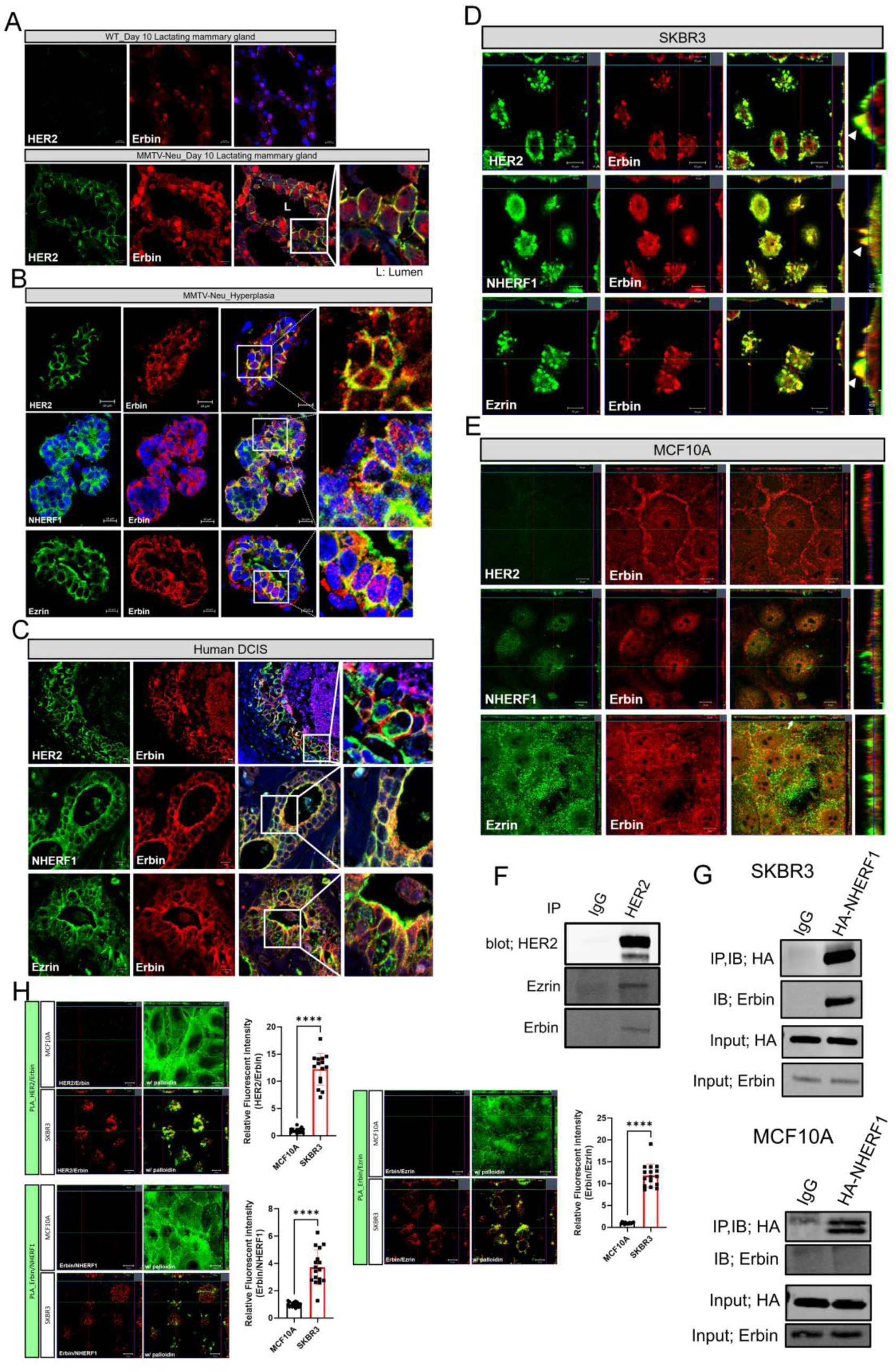
Erbin is physically associated with the HER2/NHERF1/Ezrin protein complex. A) Immunofluorescence for HER2 and Erbin in day 10 lactating normal mammary gland (top) and day 10 lactating MMTV-Neu mammary gland (bottom). All scale bars represent 10μM. B-C) Immunofluorescence for Erbin with HER2 (top row), NHERF1 (middle row), and Ezrin (bottom row) in hyperplastic lesions in MMTV-Neu mammary glands and human DCIS. All scale bars represent 10μM. D-E) Immunofluorescence for Erbin with HER2 (top row), Erbin with NHERF1 (middle row), and Erbin with Ezrin (bottom row) in SKBR3 cells (D) and MCF10A cells (E). All scale bars represent 10μM. F) Co-immunoprecipitation of HER2, Ezrin and Erbin. Extracts from SKBR3 cells were immunoprecipitated with either anti-HER2 antibody or control IgG, and blotted for HER2, Ezrin, and Erbin. G) Co-immunoprecipitation of NHERF1 and Erbin. Extracts from HA-NHERF1 over-expressing SKBR3 and MCF10A cells were immunoprecipitated with either anti-HA antibody or control IgG and blotted for HA and Erbin. H) Proximity ligation assay (PLA) showing protein interaction of HER2/Erbin, Erbin/NHERF1, and Erbin/Ezrin in MCF10A and SKBR3 cells. Phalloidin staining showed sub-cortical actin. All scale bars represent 10μM. Bar graphs show the quantification of PLA-associated fluorescence at the plasma membrane. Bar graphs represent the mean±SEM. **** denotes p<0.00005.

### Erbin is necessary for the formation of membrane protrusions and ErbB2 signaling in HER2-positive breast cancer cells

To examine the functions of Erbin in HER2 positive cells, we generated Erbin knockdown SKBR3 cells (SKBR3-ErbinKD) with a stable 85% decrease in Erbin mRNA levels (Fig. 4A). Knocking down Erbin expression decreased the levels of total and phosphorylated HER2 and EGFR proteins in SKBR3 cells, suggesting an inhibition of HER2 signaling reminiscent of inhibiting other constituents of the HER2 signaling complex within these cells. Immunofluorescence staining for HER2 demonstrated that loss of Erbin was associated with internalization of ErbB2 within vesicular appearing structures as well as the loss of NHERF1 and Ezrin co-localization with internalized ErbB2 (Fig. 4B&C). Furthermore, this was associated with a change in membrane structure. Scanning electron microscopy (SEM) images demonstrate that control SKBR3 cells have large, complex membrane protrusions as compared to MCF10A cells (Fig. 4D&E). Notably, as assessed by both SEM and immunofluorescence for phalloidin, SKBR3-ErbinKD cells showed a significant reduction in the formation of membrane protrusions (Fig. 4E-G). These findings indicate that Erbin is required for sustained HER2 signaling as well as the maintenance of protruding membrane structures previously shown to be sites of activated HER2 (6–8, 22, 23).

**Figure 4.**
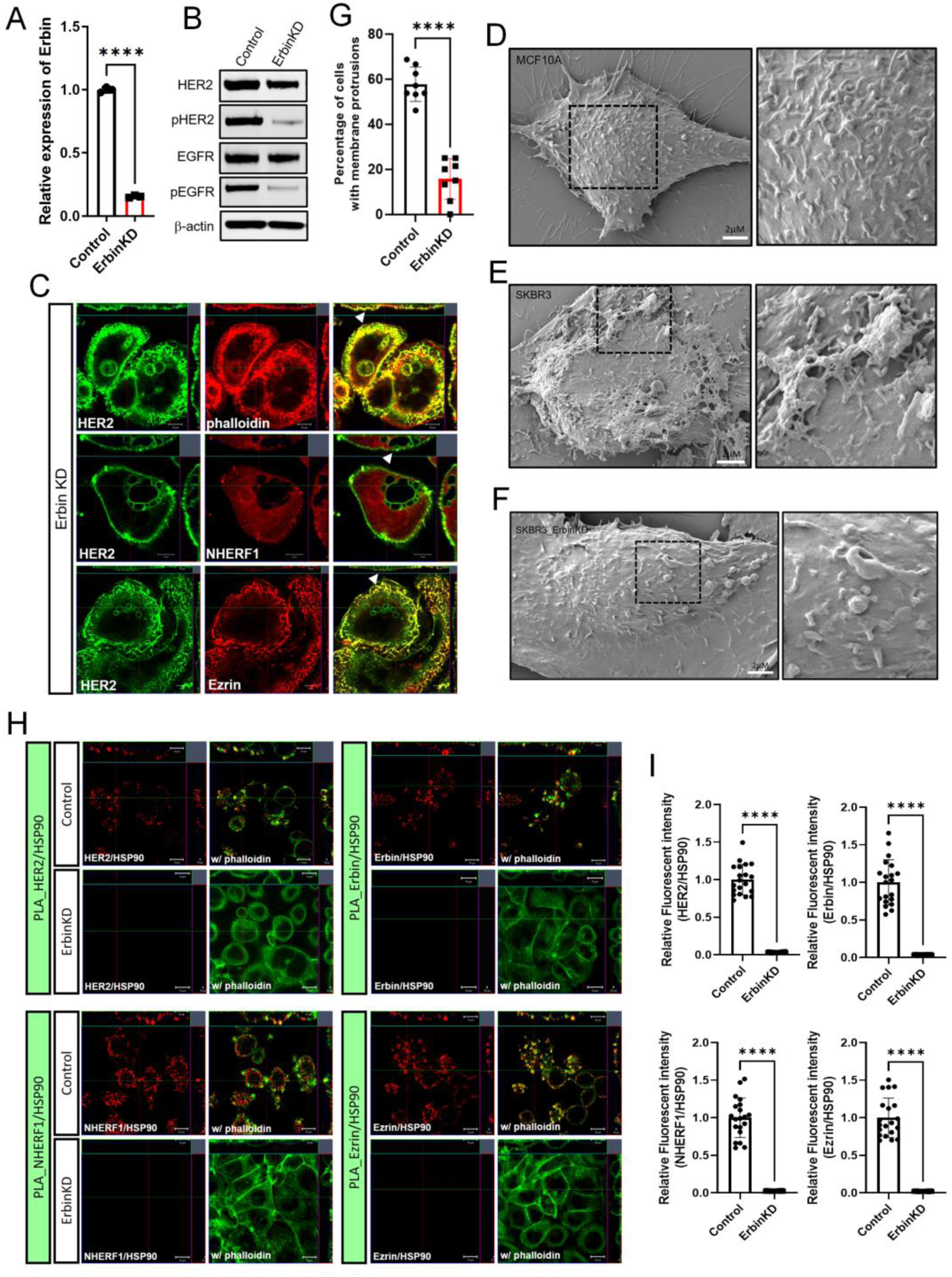
The loss of Erbin inhibits HER2 signaling and the formation of membrane protrusions. A) Erbin mRNA expression assessed by QPCR in control and Erbin knockdown SKBR3 cells. (n=3). B) Western blot analysis of HER2, phospho-HER2, EGFR, and phospho-EGFR in control and Erbin knockdown SKBR3 cells. C) Immunofluorescence for HER2 with phalloidin (top row), NHERF1 (middle row), and Ezrin (bottom row) in Erbin knock down SKBR3 cells. All scale bars represent 10μM. D-F) Scanning electron microscopy images from MCF10A, SKBR3, and Erbin knockdown SKBR3 cells. G) Percentage of cells forming membrane protrusions in control and Erbin knockdown SKBR3 cells as assessed by immunofluorescence for ____. Each data point represents the percentage of cells in 8 separate, randomly chosen microscope fields. H) PLA (red fluorescence) showing protein interaction between HER2/HSP90, Erbin/HSP90, NHERF1/HSP90, and Ezrin/HSP90 in control and Erbin knock down SKBR3 cells. Phalloidin staining in green outlines the sub-cortical actin beneath the plasma membrane. All scale bars represent 10μM. I) Quantification of PLA results (Fig. 3H) as assessed by measuring the intensity of PLA signal fluorescence. Bar graphs represent the mean±SEM. **** denotes p<0.00005.

Erbin has been reported to promote interactions between HER2 and HSP90, and HSP90 is also required to maintain activated ErbB2 within protruding membrane domains on the surface of breast cancer cells (21). Given our findings that knocking down Erbin led to loss of these structures, we also examined how it affected interactions between HER2 and HSP90. Using PLA, we found that Erbin, NHERF1, and Ezrin all interact with HSP90 within actin-rich, plasma membrane domains at the apical aspect of these cells (Fig. 4H&I). These interactions were dependent on the presence of Erbin, as evidenced by the dramatically reduction of PLA signal in SKBR3-ErbinKD cells (Fig. 4H&I). These data demonstrate that Erbin facilitates interactions of HSP90 with HER2, NHERF1 and Ezrin within plasma membrane protrusions.

### Erbin is required for interactions between HER2, NHERF1, and Ezrin within membrane protrusions

Previous studies have identified interactions between HER2, NHERF1 and Ezrin that are important to maintain HER2 within membrane protrusions at the cell surface (6–8). Using SKBR3-ErbinKD cells, we next performed PLA to test whether Erbin is involved in the formation of interactions between Ezrin, NHERF1 and HER2 in SKBR3 cells. First, no PLA signal was seen in non-transformed MCF10A. In contrast, there were clear interactions between HER2 and NHERF1, HER2 and Ezrin, and Ezrin and NHERF1 primarily at the cell surface and concentrated within membrane protrusions in control SKBR3 cells (Fig. 5A-C). Loss of Erbin did not eliminate physical interactions between HER2 and NHERF1 but shifted them away from membrane structures towards a more intracellular and basolateral distribution (Fig. 5A). Interestingly, there was not a significant change in the PLA signal for HER2 and Ezrin although these interactions appeared more diffusely within the apical membrane and were also increased intracellularly (Fig. 5B). Finally, Erbin knockdown dramatically decreased interactions between of NHERF1 and Ezrin overall and specifically within membrane protrusions (Fig. 5C). These data suggest that while Erbin may not be required for HER2 to interact with NHERF1 or Ezrin individually, it is important for facilitating NHERF1 and Ezrin interactions that maintain HER2 within actin-rich, protruding membrane domains within SKBR3 cells that past work has shown to be important areas of active HER2 signaling.

**Figure 5.**
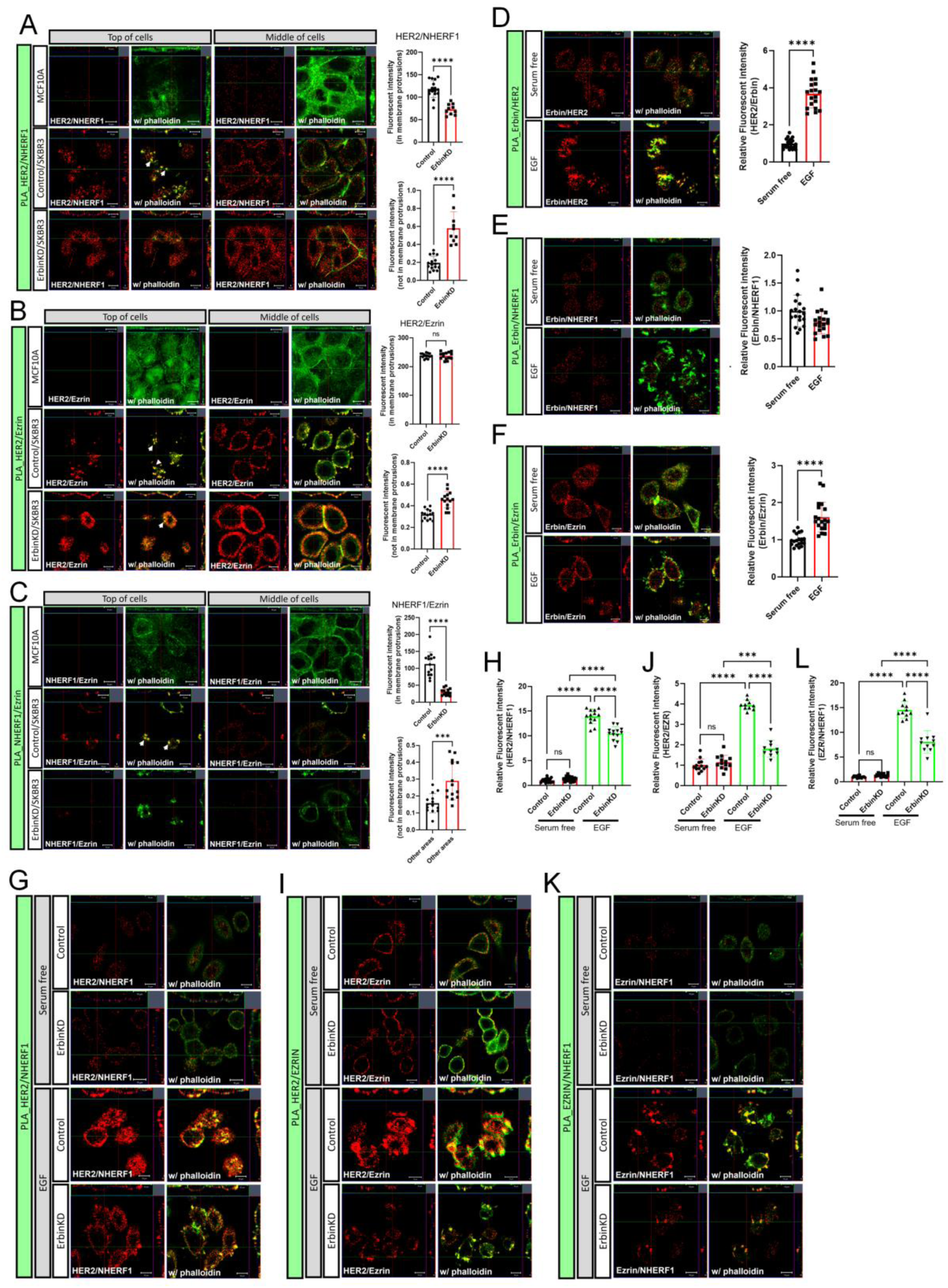
Erbin is required for interactions between HER2/NHERF1/Ezrin in membrane protrusions. A-C) PLA fluorescence (red) showing protein interaction between HER2/NHERF1 (A), HER2/Ezrin (B), and NHERF1/Ezrin (C) in MCF10A, SKBR3, and Erbin knockdown SKBR3 cells. Top of cells; focused on surface of plasma membrane, Middle of cells; focused on the mid-portion of cells to examine intracellular interactions. Phalloidin staining in green outlines the sub-cortical actin beneath the plasma membrane. All scale bars represent 10μM. Bar graphs show quantification of PLA results by measuring intensity of fluorescence from PLA signal at the surface and inside of cells. Bars represent the mean±SEM. D-F) PLA showing protein interaction of HER2/Erbin (D), Erbin/NHERF1 (E), and Erbin/Ezrin (F) in serum free (SF) and EGF treated SKBR3 cells. Phalloidin staining is in green. All scale bars represent 10μM. Bar graphs show quantification of PLA results by measuring intensity of fluorescence from PLA signal. G-L) PLA showing protein interaction between HER2/NHERF1 (G), HER2/Ezrin (I), and NHERF1/Ezrin (K) in control and Erbin knockdown SKBR3 cells under either serum free or serum free with 100ng/ml EGF. All scale bars represent 10μM. H, J, and L) Bar graphs show quantification of PLA results by measuring intensity of fluorescence from PLA signal at the cell membrane. Bar graphs represent the mean±SEM. *** denotes p<0.0005, **** denotes p<0.00005.

### Erbin regulates EGF-stimulated formation of an HER2/NHERF1/Ezrin complex

Activation of HER2 by EGF facilitates the formation of a multi-protein complex that includes HER2, NHERF1, and Ezrin, and also promotes the formation of larger and more complex membrane protrusions (6–8, 22). Therefore, we next queried whether Erbin is involved in the responses to EGF. We used PLA to assess changes in the protein-protein interactions between Erbin, HER2, NHERF1 and Ezrin in response to EGF. EGF treatment resulted in a significant increase in the physical interactions between Erbin and HER2 as well as Erbin and Ezrin (Fig. 5D&F). However, interactions between Erbin and NHERF1 were unchanged by EGF (Fig. 5E). We also examined protein-protein interactions and their response to EGF by PLA using SKBR3-ErbinKD cells. First, knocking down Erbin expression did not change the low level of PLA signals seen in serum starved cells for interactions between HER2/NHERF1, HER2/Ezrin or Ezrin/NHERF1. On the other hand, EGF treatment dramatically stimulated PLA signals indicating interactions between HER2 and NHERF1, HER2 and ezrin and ezrin and NHERF1 in the control SKBR3 cells (Fig. 5G-L). However, these interactions were significantly reduced by knocking down Erbin expression in SKBR3 cells (Fig. 5G-L). As before, interactions between NHERF1 and HER2 were somewhat less sensitive to loss of Erbin than were interactions between HER2 and Ezrin or ezrin and NHERF1.

### NHERF1 and Ezrin are required for Erbin to interact with HER2

Next, we tested whether NHERF1 and Ezrin are required for Erbin to interact with HER2. NSC668394 is an Ezrin inhibitor that disrupts the formation of membrane protrusions and leads to the internalization and degradation of HER2, NHERF1, and Ezrin (8). Likewise, we have reported that the loss of NHERF1 expression has similar effects on HER2 localization and signaling (7). Both NSC668394 treatment and NHERF1 knockdown in SKBR3 cells (SKBR3-NHERF1KD) resulted in loss of co-localization of Erbin with HER2, NHERF1 or Ezrin (Fig. 6A&B). Furthermore, HER2, NHERF1 and Ezrin could be detected internalized away from the plasma membrane (Fig. 6A&B). PLA further confirmed that treatment with NSC668394 or NHERF1 knockdown caused significant reductions in the physical association of Erbin with HER2, NHERF1 and Ezrin (Fig. 6C-D). These results demonstrate that Ezrin and NHERF1 are required to facilitate interactions between HER2 and Erbin.

**Figure 6.**
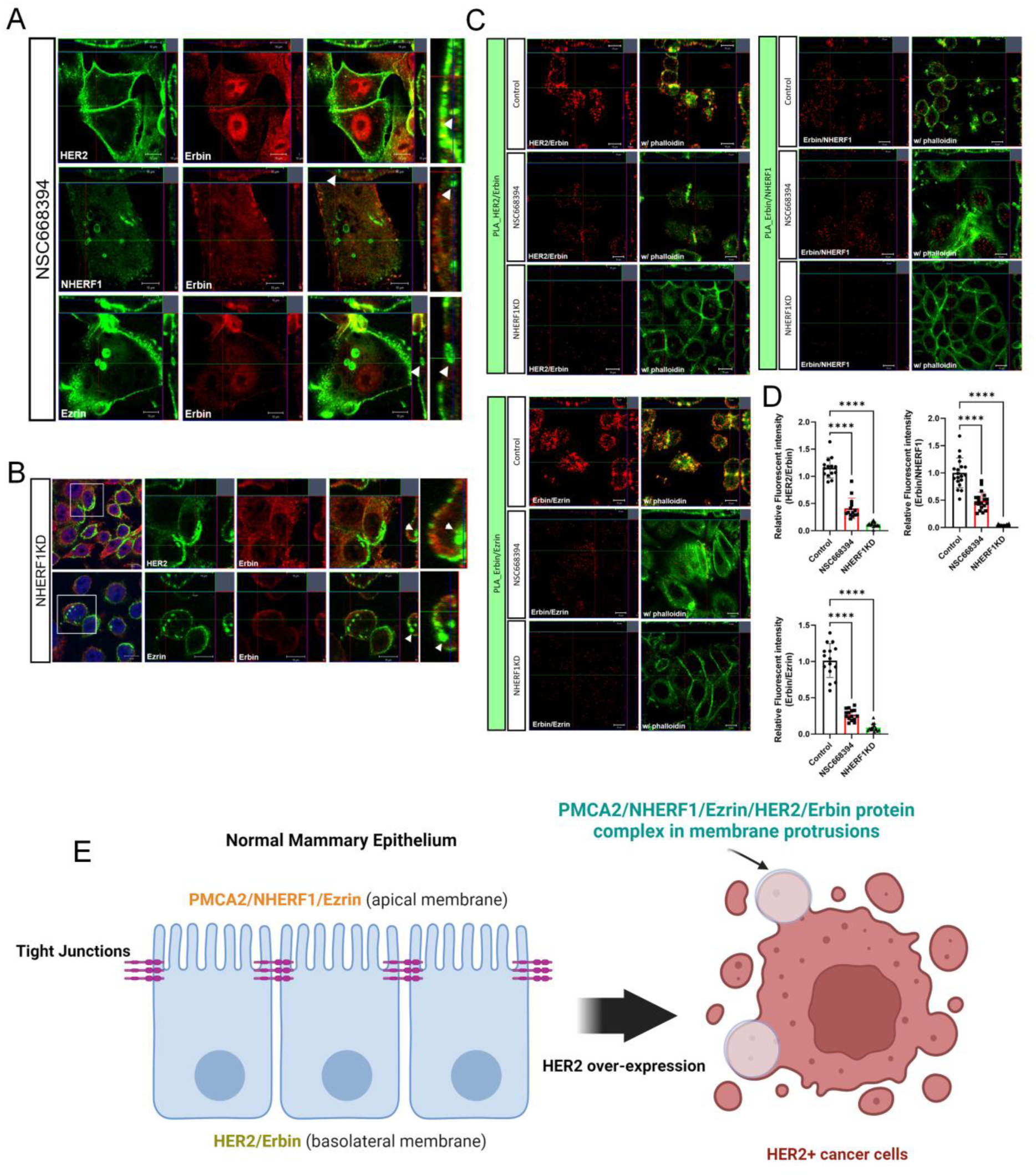
Ezrin and NHERF1 are required for Erbin and HER2 interactions. A) Immunofluorescence for Erbin with HER2 (top row), NHERF1 (middle row), and Ezrin (bottom row) in NSC668394 treated SKBR3 cells. All scale bars represent 10μM. B) Immunofluorescence for Erbin with HER2 (top row) and Ezrin (bottom row) in NHERF1 knockdown SKBR3 cells. All scale bars represent 10μM. C) PLA fluourescence showing protein-protein interactions between HER2/Erbin, Erbin/NHERF1, or Erbin/Ezrin in control, NSC668394 treated-, and NHERF1 knock down SKBR3 cells. All scale bars represent 10μM. D) Quantification of PLA results was performed by measuring the fluorescent intensity of PLA signals at the cell surface. Bar graphs represent the mean±SEM. **** denotes p<0.00005. E) A working model proposing how HER2 mediated disruption of apical/basal polarity creates the opportunity for the formation of a multi-protein signaling complex that includes HER2, Erbin, PMCA2, NHERF1, and Ezrin. Created with BioRender.com.

## Discussion

Erbin is a member of the leucine-rich repeat and PDZ (PSD95/DLG/ZO-1) domain (LAP) protein family. LAP proteins localize to the basolateral membrane, physically associate with junctional proteins and are involved in organizing and maintaining basolateral polarity in epithelial cells (13, 16, 24). Ezrin and NHERF1 are scaffolding proteins that localize to the apical membrane of polarized cells and function to maintain apical polarity, to stabilize apically located membrane proteins to the actin cytoskeleton and to organize and maintain apical membrane structures such as microvilli and stereocilia (17, 25–27). Given their polarized localization in epithelial cells, it is unlikely that Erbin interacts with either Ezrin or NHERF1 in normal cells (Fig. 6E). Therefore, we found it interesting that inhibiting Erbin reduced HER2 expression and signaling within breast cancer cells and reduced the formation and growth of HER2-dependent mammary tumors in MMTV-Neu mice in a similar fashion to what we have previously shown for inhibition of Ezrin and NHERF1. Therefore, we hypothesized that Erbin might participate in the same NHERF1/Ezrin/HER2 complex that we have previously described in HER2-positive breast cancer cells (Fig. 6E). Indeed, in this report we demonstrate this to be the case. We show that Erbin co-localizes with and is in close physical proximity to HER2, NHERF1 and Ezrin in breast cancer cells, especially within actin-rich membrane protrusions. However, these proteins do not co-localize in normal murine mammary epithelium or in non-malignant MCF10A human breast epithelial cells. Furthermore, NHERF1 and Ezrin are required for Erbin to interact with HER2 and Erbin is required for NHERF1 to interact with HER2 or Ezrin. Interestingly, knocking down Erbin did not affect interactions between HER2 and ezrin. Finally, knocking down Erbin expression led to the effacement of the complex plasma membrane protrusions evident in SKBR3 cells and led to the internalization of HER2 away from the plasma membrane. Thus, in malignant cells, Erbin becomes associated with HER2, Ezrin, NHERF1 and HSP90, and this multiprotein complex leads to the stabilization of activated HER2 at the plasma membrane within specific signaling domains that form actin-rich membrane protrusions.

It is instructive to compare the expression pattern of Ezrin, NHERF1, Erbin and HER2 by immunofluorescence in the mouse mammary gland. In normal mice, we did not detect any appreciable HER2 expression while NHERF1 and ezrin were expressed in the apical membrane, Erbin was expressed predominantly in the cytoplasm and Erbin did not co-localize with either Ezrin or NHERF1. In MMTV-Neu mice, Erbin could be found within the basolateral membrane of non-transformed mammary epithelial cells, where it co-localized with HER2. In non-transformed cells, Ezrin and NHERF1 expression remained apical and did not co-localize with HER2. However, in areas of hyperplasia or invasive tumors in MMTV-Neu mice, HER2, Erbin, NHERF1 and Ezrin were located more diffusely within the plasma membrane and all co-localized with each other. Interestingly, we identified rare areas of apparently normal appearing epithelium adjacent to human DCIS lesions, in which HER2 remained co-localized with Na/K ATPase in the basolateral membrane, but ezrin and NHERF1 expression spread from the apical membrane into the basolateral compartment. However, more quantitative observations will be required to determine if this is a common transitional pattern between normal cells and DCIS lesions.

Like other epithelial cells, luminal epithelial cells in the breast are polarized with respect to their apical (lumen facing) and basolateral surfaces. NHERF1 and Ezrin are both localized specifically at the apical plasma membrane of mammary and other epithelial cells, and participate in the formation and maintenance of apical-basal polarity (25–28). In addition, both NHERF1 and Ezrin play critical roles in the organization of microvilli (25, 29, 30). NHERF1 stabilizes transmembrane proteins to the apical membrane through its PDZ domain and its ability to bind Ezrin and other ERM proteins, which interact directly with the cytoskeleton. The separation between apical and lateral membrane domains is actively controlled by the Par complex, which is associated with tight junctions that form a diffusion barrier separating apical and lateral membrane proteins from each other as well as controlling diffusion of ions and fluids between epithelial cells. The Par complex consists of a series of intracellular proteins, including Par3, Par6, cdc42 or rac2, and atypical PKC, which associate with tight junction proteins through direct interactions between junctional proteins and Par3. Aranda and colleagues showed that activated ErbB2 could disrupt this complex by associating with Par6 and aPKC at the expense of their interactions with Par3. In MDCK kidney epithelial cells and in MCF10A mammary epithelial cells, this resulted in disruption of tight junctions and diffusion of apical proteins into the basolateral membrane. This model is consistent with our observations that in relatively normal-appearing cells at the borders of human DCIS lesions, we see continued basolateral expression of HER2 while Ezrin and NHERF1 expression break the apical-lateral border and extend into the basolateral compartment to co-localize with HER2. We confirm that HER2 interacts with Erbin at the basolateral surface in normal appearing cells in MMTV-Neu mice that overexpress HER2. By the time we observe lumen filling lesions, or in HER2-overexpressing breast cancer cells, basolateral polarization is also lost and HER2, Ezrin, NHERF1, HSP90, and Erbin all join together to form stable signaling complexes that localize within protruding membrane domains on SKBR3 cells. As a result, we propose a working model that explains the formation of this multi-protein HER2 signaling platform in some developing HER2+ tumors: 1) initial upregulation of HER2 expression, 2) HER2 association with Erbin at the basolateral membrane, 3) HER2-mediated disruption of apical/basal polarity, 4) lateral diffusion of NHERF1, Ezrin, PMCA2 and other proteins, which, in turn form novel physical interactions with HER2/Erbin complexes, 5) enhanced HER2 membrane stability and signaling, and 6) tumor progression.

In summary, in breast cancer cells, Erbin interacts with HER2 as well as other scaffolding proteins, including NHERF1 and Ezrin. Our results suggest that protein-protein interactions between Erbin, Ezrin and NHERF1 may be important for HER2-driven tumorigenesis. The loss of Erbin in SKBR3 cells disrupts multiple other protein-protein interactions that support HER2 signaling platforms. It also disrupts membrane architecture, HER2 membrane stability and HER2 biochemical signaling. Future investigations to identify how these different proteins come together and interact during HER2-driven transformation may help in the development of new prognostic and therapeutic tools to target HER2-positive breast cancers.

## Materials and methods

### Materials

We obtained the Ezrin inhibitor (NSC668394) from Calbiochem.

### Cell culture

SKBR3 cells were obtained from ATCC and maintained in culture in DMEM +GlutaMAX-1 (Gibco-life Technologies) containing 10% fetal bovine serum (FBS) and pen/strep (Gibco-life Technologies) at 37°C in 5% CO_2_. In some experiments, cells were cultured as above but in media without FBS for 16 hours and then were treated with 100ng/ml EGF (Cell Signaling) for 2 hours. In other experiments, cells were treated with NSC668394 at 10μM for 16 hours. MCF10A cells were cultured in DMEM/F12 (Gibco-Life Technologies) containing 5% horse serum, EGF (100ng/ml), hydrocortisone (1mg/ml), cholera toxin (1mg/ml), insulin (10μg/ml), and pen/strep (Gibco-Life Technologies) at 37°C in 5% CO^2^.

### Erbin knockdown cell line

A stable cell line expressing shRNA directed against Erbin was generated by transducing cells with commercially prepared lentiviruses containing three individual shRNA targetting Erbin mRNA (sc-40541-V) (Santa Cruz). Cells were cultured in 6-well plates and infected by adding the shRNA lentiviral particles to the culture for 48 hours per the manufacturer’s instructions. Stable clones expressing the specific shRNAs were selected using 5μg/ml of puromycin (Gibco-life technologies) and were pooled together to generate the cells used in the experiments. NHERF1 knock-down SKBR3 cells were generated previously in a similar fashion (7).

### Spatial transcriptomics data analysis

The Visium dataset used in this analysis was provided by 10X Genomics. Data for human breast cancer sections are freely available and can be downloaded from the 10X website (https://www.10xgenomics.com/resources/datasets). Gene expression from the visium data was normalized and high-variance features were identified using the SCTransform() function from the R package ‘Seurat’ (31) (version 4.4.0). Visium spots were annotated using the R package ‘Cottrazm’ (32) (version 0.1.1), which distinguishes malignant spots from boundary and non-malignant spots based on their copy number variations, and cell type annotations provided by a previous report (19). Spots from ‘DCIS’ were classified into high-expressing and low-expressing groups based on the median expression values of SLC9A3R1, EZR, and ERBIN. Differentially expressed genes (DEGs) between the two groups were identified using the FindMarkers() function in ‘Seurat’ (version 4.4.0). Gene Set Enrichment Analysis was then performed using the DEGs and hallmark gene sets from the Human MSigDB Collections (version 2023.1) with the R package ‘fgsea’ (version 1.25.1).

### Public RNA sequencing data analysis

Preprocessed data matrices from published RNA sequencing datasets were downloaded from the NCBI Gene Expresison Omnibus (GSE69240) (33). The log2CPM values were inversely transformed to count value then differential expression of HER2_high human DCIS and normal breast tissues was performed using DESeq2 package (version 3.16). Normal(n=10) and ERBB2_high(n=13) data were grouped. Gene Set Enrichment Analysis of differentially expressed genes was determined using the clusterProfiler package (version 4.4) from the molecular signature database Hallmark collection (version 7.4) (34).

### Immunofluorescence

Cells were grown on coverslips, fixed in 4% paraformaldehyde for 20 min, permeabilized with 0.2 % Triton X100 for 10 mins, washed 3 times with PBS and incubated with primary antibody overnight at 4°C. The cells were washed 3 times with PBS and incubated with secondary antibody for 1 hour at room temperature. After washing, coverslips were mounted using Prolong Gold antifade reagent with DAPI (Invitrogen). Paraffin-embedded tissue sections were cleared with histoclear (National Diagnostics) and graded alcohol using standard techniques. Antigen retrieval was performed using 7mM citrate buffer, pH 6.0 under pressure. Sections were incubated with primary antibody overnight at 4°C and with secondary antibody for 1 hour at room temperature. Coverslips were mounted using Prolong Gold antifade reagent with DAPI (Invitrogen). All images were obtained using a Zeiss 780 confocal microscope and Zeiss LSM 880, and settings were adjusted to allow for detection of fine membrane structure. Primary antibodies were against: NHERF1 (sc134485, sc271552), Na/K ATPase (sc28800), Ezrin (sc58758) from Santa Cruz (Dallas, TX); Ezrin (3145) from cell signaling (Danvers, MA); HER2 (MA1-35720) from Invitrogen (Grand Island, NY); Erbin (NBP256104) from NOVUS (Centennial, CO).

### Immunoblotting

Protein extracts were subjected to SDS-PAGE and transferred to a nitrocellulose membrane by wet western blot transfer (Bio-Rad). The membrane was blocked in TBST buffer (TBS + 1% Tween) containing 5% milk for 1 hour at room temperature. The blocked membranes were incubated overnight at 4 °C with primary antibodies in Odyssey blocking buffer, 927-40000, washed 3 times with TBST buffer, and then incubated with secondary antibodies provided by LI-COR for 2 hours at room temperature. After 3 washes with TBST buffer, the membranes were analyzed using the ODYSSEY Infrared Imaging system (LI-COR). Primary antibodies were against: HER2 (sc33684), mouse β-actin (sc-69879) from Santa Cruz (Dallas, TX) from Santa Cruz (Dallas, TX); EGFR (4267), phosphor-EGFR (2234), phosphor-HER2 (2247) from cell signaling (Danvers, MA);. All immunoblot experiments were performed at least 3 times and representative blots are shown in the figures.

### Co-immunoprecipitation

Cells were lysed with RIPA buffer (1% NP-40, 0.5% sodium deoxycholate, 0.1% SDS, 20mM Tris Hcl, and 150mM NaCl), and cell extracts were incubated overnight at 4°C with protein A/G beads (sc-2003, Santa Cruz) and the specific antibody. After centrifugation, the immunoprecipitated proteins were eluted with LDS sample buffer containing 10% beta-mercaptoethanol. The resulting samples were then analyzed by Western blot.

### Proximity Ligation Assays

Proximity ligation assays (PLA) were performed using the Duolink™ assay kit (Sigma). SkBR3 and MCF10A cells were seeded onto 22 mm round collagen-coated coverslips (Corning, Cat N 354089). The experiments were performed when the cells reached 80% confluence. The cells were washed 3 times with PBS and paraformaldehyde (4% in PBS) was added to each well for 20 min, after which they were permeabilized with 0.2% Triton X100 in PBS. Permeabilized cells were incubated with combinations of the following antibodies: HER2 (MA1-35720), Erbin (NBP2-13968), NHERF1 (sc-134485 and sc271552), Ezrin (3145 and sc58758), and HSP90 (ab203085). PLA probes were then added, and the assay was performed as per the manufacturer’s instructions. All images were obtained using a Zeiss 780 confocal microscope and Zeiss LSM 880, and settings were adjusted to allow for detection of fine membrane structure. Quantification was done by measuring the intensity of fluorescence at randomly chosen phalloidin positive area at the plasma membrane.

### RNA Extraction and Real-Time RT-PCR

RNA was isolated using TRIzol (Invitrogen). Quantitative RT-PCR was performed with the SuperScript III Platinum One-Step qRT-PCR Kit (Invitrogen) using a Step One Plus Real-Time PCR System (Applied Biosystems) and the following TaqMan primer sets: human Erbin (Hs1049966_m1). Human HPRT1 (4325801) was used as reference genes (Invitrogen). Relative mRNA expression was determined using the Step One Software v2.2.2 (Applied Biosystems).

### Statistics

Statistical analyses were performed with Prism 7.0 (GraphPad Software, La Jolla, CA). Statistical significance was determined by using unpaired t test for comparisons between 2 groups and one-way ANOVA for groups of 3 or more. All bar graphs represent the mean±SEM, * denotes p<0.05, ** denotes p<0.005, *** denotes p<0.0005, **** denotes p<0.00005.

## Acknowledgments

NRF2022R1A4A2000827 from the National Research Foundation of Korea (NRF) to J.Choi. R01 HD100468 and R01 HD076248 from the NIH to J. Wysolmerski.

## Conflict of interest

Nothing to declare.

## Notes

### Competing Interest Statement

The authors have declared no competing interest.

